# The human PDZome 2.0: characterization of a new resource to test for PDZ interactions by Yeast Two-Hybrid

**DOI:** 10.1101/2020.08.06.239343

**Authors:** Monica Castro-Cruz, Frédérique Lembo, Jean-Paul Borg, Gilles Trave, Renaud Vincentelli, Pascale Zimmermann

## Abstract

PSD95-disc large-zonula occludens (PDZ) domains are globular modules of 80-90 amino acids that co-evolved with multicellularity. They commonly bind to carboxy-terminal sequences of a plethora of membrane-associated proteins and influence their trafficking and signaling. We previously built a PDZ resource (PDZome) allowing to unveil human PDZ interactions by Yeast two-hybrid. Yet, this resource is partial according to the current knowledge on the human PDZ proteome. Here we built the PDZome 2.0 library for Yeast two-hybrid, based in a PDZ library manually curated from online resources. The PDZome2.0 contains 305 individual clones (266 PDZ domains in isolation and 39 tandems), for which all boundaries have been designed based on available PDZ structures. Using as bait the E6 oncoprotein from HPV16, a known promiscuous PDZ interactor, we show that PDZome 2.0 outperforms the previous resource.

## Introduction

PDZ scaffold proteins are involved in a wide range of cellular processes including the establishment and maintenance of polarity, protein trafficking, signaling and the coordination of synaptic events (1–3). They contain one or more PDZ domain that is an abundant and promiscuous protein interaction module. PDZ domains were first identified in the proteins **P**SD-95 (postsynaptic density-95), **D**lg-1 (disc large-1), and **Z**O-1 (zona occludens-1) (4–8). PDZ domains generally recognize short linear motifs of minimum 4 amino acids (PDZ binding motifs or PBM) located at the C-terminal region of receptors, co-receptors, or adhesion molecules (9). Additionally, PDZ domains can interact with internal protein motifs, lipids and other PDZ domains (10–12). PDZ interactions can be tuned in various ways. Changes in salt content and pH (13), auto-inhibition (14), allosteric regulation (15) and phosphorylation (16) are some of the features that modulate PDZ interactions (for reviews see (17,18)).

PDZ domains are composed by 80-90 amino acid residues which fold in six β-strands (A-F) and two α-helices (A-B), forming a partially opened antiparallel B barrel structure (1,19). The PBM binds in a groove formed by the α-helix B and the β-sheet B (19). The PDZ binding groove is connected by a loop which often contains the GLGF motif. The GLGF motif, also described as R/K-X-X-X-G-φ-G- motif where X is any and φ is an hydrophobic residue, can vary significantly and contributes to the affinity of the interactions with the PBM (19,20). Structural and functional studies suggest that PDZ domains prefer specific residues in a PBM. One can currently identify three main PBM classes that can occur in 16 specificity sub-classes (20). Yet, approaches like e.g. phage display suggest that PBM specifities go beyond such classification (21,22).

PDZ domains are rare in non-metazoans. For example, bacteria and yeast display no more than 2 and 4 PDZ-domain containing proteins, respectively (23,24). In contrast, PDZ proteins are abundant in metazoans, suggesting they co-evolved with multicellularity (24). Several studies based on sequence analysis using SMART (http://www.smart.embl-heidelberg.de), Interpro (https://www.ebi.ac.uk/interpro/), and PFAM (https://pfam.xfam.org/) suggest that the number of PDZ domains in the human proteome ranges from 234 to 450 (25–27). Based on these strictly *in silico* studies, a first collection of human PDZ domains was built (PDZome) to test for PDZ interactions by Yeast-two-hybrid (Y2H) (27). This resource contains 246 PDZ domains. Yet according to more refined study including a 3D-structure based approach and careful manual annotation, this resource contains PDZ domains truncated at their N- and C-termini, by 5 to 16 amino-acids (28). Such truncation might compromise proper folding and binding activities (28–31). In this refined study, we identified 266 PDZ domains embedded in 150 proteins (omitting spliced forms) in the human proteome.

Noteworthy, it became clear that some PDZ domains occurring in tandem (separated by a short conserved linker region) can function as supramodules (32,33). The binding properties of these supramodules are different from those of PDZ domains taken in isolation. Generally, PDZ tandems display higher affinity for their target and in some case the tandem might be necessary for proper folding of individual domains (33,34).

Because the original Y2H PDZome resource (27) misses some PDZ domains, does not contain tandems and also because of the presence of suboptimal boundaries, we prepared a new resource that we called PDZome 2.0. The PDZome 2.0, is more comprehensive including the 266 manually annotated sequences of single PDZ domains (28). Additionally, it contains 39 PDZ domains in tandem. To test for the performance of PDZome 2.0, we used the E6 oncoprotein present in the human papilloma virus-16 (HPV16). The PDZome 2.0 detected a total of 54 E6-PDZ interactions. Twenty-nine are common with the 36 previously identified by the PDZome and 25 are newly identified. We therefore propose the PDZome 2.0 as a more performant resource to comprehensively map human PDZ interactions by Y2H approach.

## Materials and Methods

### Sub-cloning of prey and baits

Prey entry clones were collected in the pZeo or the pDONOR201 Gateway ® vectors (NZYTech, Ltd.). All the entry clones were subcloned into the Y2H expression vector pACT2-AD using Gateway ® LR reactions (Invitrogen). After sequence validation, all pACT2-AD clones were transformed into the haploid Y187 yeast strain (MATα, ura3-52, his3-200, ade2-101, trp1-901, leu2-3, 112, gal4Δ, met-, gal80Δ, MEL1, URA3::GAL1UAS - GAL1TATA-lacZ).

The two baits used here, correspond to a fragment of the HVP16 E6 oncoprotein wildtype (MSCCRSSRTRRETQL), and the same fragment without the PDZ binding motif or ΔTQL (MSCCRSSRTRRE). The E6 fragments were subcloned into the pGBT9-BD vector for expression in yeast, as reported previously (27). After sequence validation, E6 constructs were transformed into the haploid AH109 yeast strain (MATa, trp1-901, leu2-3, 112, ura3-52, his3-200, gal4Δ, gal80Δ, LYS2::GAL1UAS -GAL1TATA -HIS3, GAL2UAS -GAL2TATA -ADE2, URA3::MEL1UAS -MEL1 TATA -lacZ).

### Y2H assays

PDZ interactions were tested screened by Y2H assay (36). Briefly, the Y2H was performed through mating of the two yeast strains Y187 (α) and AH109 (a). The yeasts were grown together (α + a) in liquid Yeast extract-Peptone-Dextrose (YPD) supplemented with 10% PEG for 5 - 6 h at 30 °C under gentle agitation (140 rpm). After one wash in sterile water, the yeasts were spotted on solid medium. To test the mating efficiency, the yeasts were spotted on a solid permissive medium SC Agar -L -W. To test for interactions, the yeasts were spotted on a solid selective medium SC Agar -L -W -H. All SC-Agar plates were incubated at least 72 h and up to 1 week at 30 °C or 2 weeks at room temperature. Images from the solid selective medium plates were captured and analyzed. Random positive clones were verified by PCR amplification and automated sequencing with the GAL-AD primer (Eurofins GATC).

## Results

### Construction of the human PDZ resource for Y2H assays

To build the human PDZome 2.0 resource allowing to test for PDZ interactions by Y2H, the 266 known human PDZ domain sequences (**Table S1**), bearing boundaries optimized based on available structural data (28), were introduced in the prey vector by Gateway ® approach (Fig. 1A). We also included 39 PDZ tandems (**Table S2**). The PDZ tandems were designed using the online UniProt resource (https://www.uniprot.org/). First, all PDZ proteins with more than one PDZ domain were included in the list (multi-PDZ proteins). Then, within these multi-PDZ proteins, those in which 2 PDZ domains were connected by a linker region of up to 36 amino acid residues acids were included. The final list of 39 tandems, belonging to 28 PDZ proteins, represent around 20% of the human PDZ proteome (**Table S2**).

**Fig 1.**
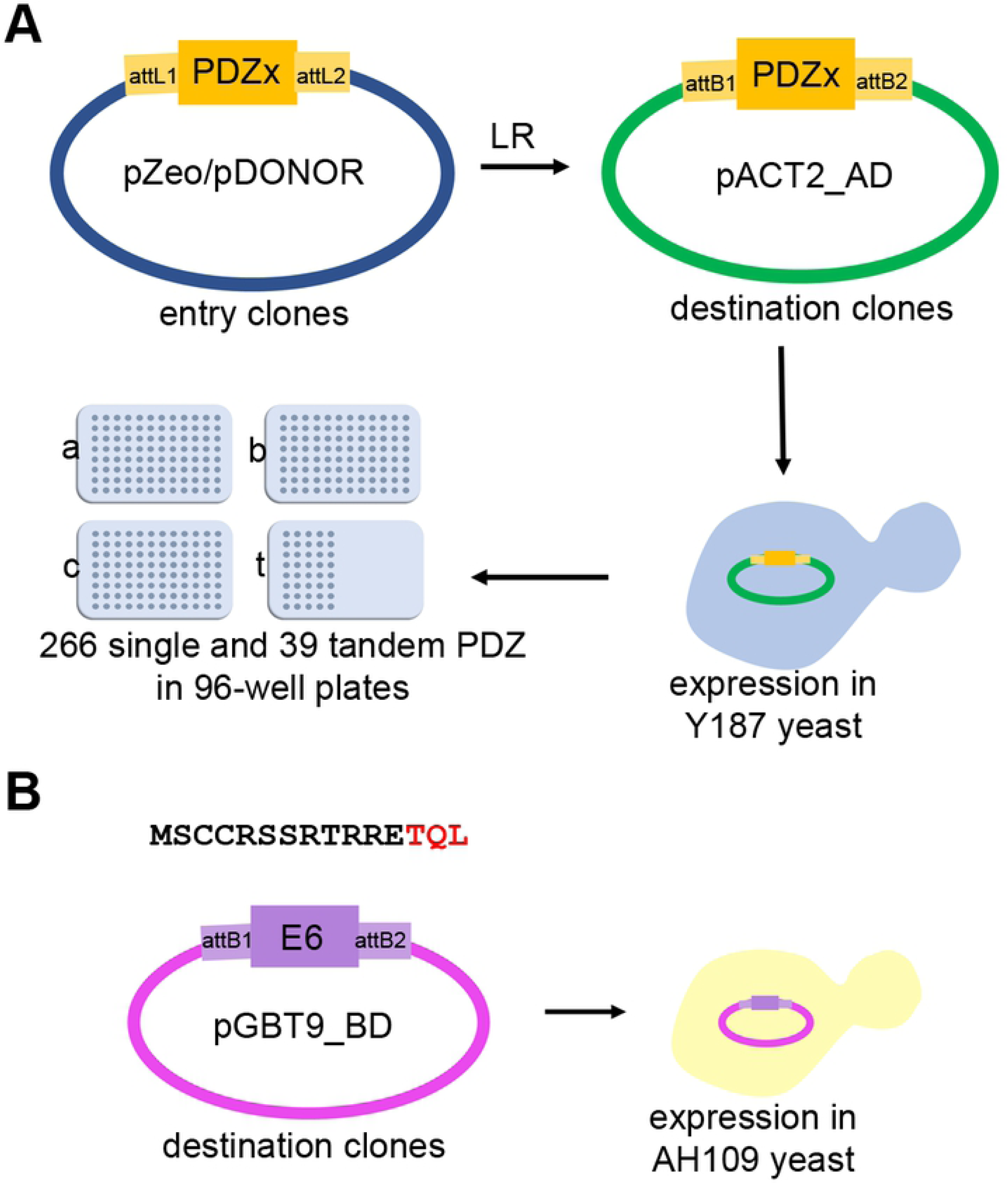
Construction of the PDZome 2.0 for yeast two-hybrid screens. The PDZome 2.0 was built using the Gateway ® cloning system. (**A**) The entry clones corresponding to the open reading frames (ORF) of the 266 PDZ domains and 39 PDZ domains in tandem were subcloned from pZeo or pDONOR entry vectors. The ORFs were then introduced into the pACT2-AD vector using Gateway ® LR clonase. After validation by sequencing, pACT2-AD clones were transformed into the Y187 (type α) yeast strain. Ready for mating yeast containing the PDZome fused to the Gal4 activation domain were arranged in 4 plates of 96 wells (a, b, c correspond to single PDZ domains, whereas t corresponds to tandems). (**B**) Two peptides corresponding to the C-terminal part of the E6 protein from the HPV16 were used as baits. The wild type (MSCCRSSRTRRETQL) and the ΔTQL (or ΔPBM) were subcloned in the pGBT9-BD vector as described previously (27) and transformed into the AH109 (type a) yeast strain.

All recombinant clones present in the prey pACT2-AD vector were transformed into the haploid Y187 (α) yeast strain. The final collection of individual clones was arrayed in four 96-well plates (Fig. 1 A).

### The PDZome 2.0 for Y2H screenings is validated using the HPV16 E6 oncoprotein

To characterize the performance of the PDZome 2.0 we used a fragment of the HPV16 E6 oncoprotein as bait in Y2H screenings. The HVP16 E6 oncoprotein is involved in the development of human cervical cancer by exploiting its class I PBM which has been previously described to bind at least 29 PDZ scaffold proteins (27,37–39).

Two E6 constructs were used to validate the new resource. The wild-type E6 (MSCCRSSRTRRETQL) and the mutant E6 ΔPBM, in which the PBM is disrupted by removing the last 3 amino acids (MSCCRSSRTRRE) (Fig. 1 B). Bait constructs were subcloned in the pGBT9-BD expression vector and fusion proteins were expressed in the AH109 yeast strain for Y2H (Fig. 1 B).

Y2H screens were carried out by mating the two recombinant yeast strains Y187 (α) and AH109 (a), allowing the formation of diploid yeasts expressing the prey and the bait constructs (Fig. 2 A). According to Y2H principles, in case the E6 bait interacts with a given PDZ prey, a complex is formed and the activating domain (AD) is recruited near the reporter gene, where it can stimulate its expression (Fig. 2 B, C). To control mating efficiency, we cultured our mated yeasts in SC-Agar medium lacking leucine and tryptophan (-LW). The growth of dense white colonies indicated an efficient mating (Fig. 2 C **upper panel**). Simultaneously, to test for PDZ interactions, the mated yeasts were grown in SC-Agar medium lacking leucine, tryptophan, and histidine (-LWH). The growth of dense white colonies in the medium -LWH were indicative of E6-PDZ interactions (Fig. 2 C **middle panel**). As expected, when the E6 PBM was disrupted (E6 ΔTQL), yeasts failed to grow in the -LWH medium (Fig. 2 C **lower panel**) indicating that the PBM is essential.

**Fig 2.**
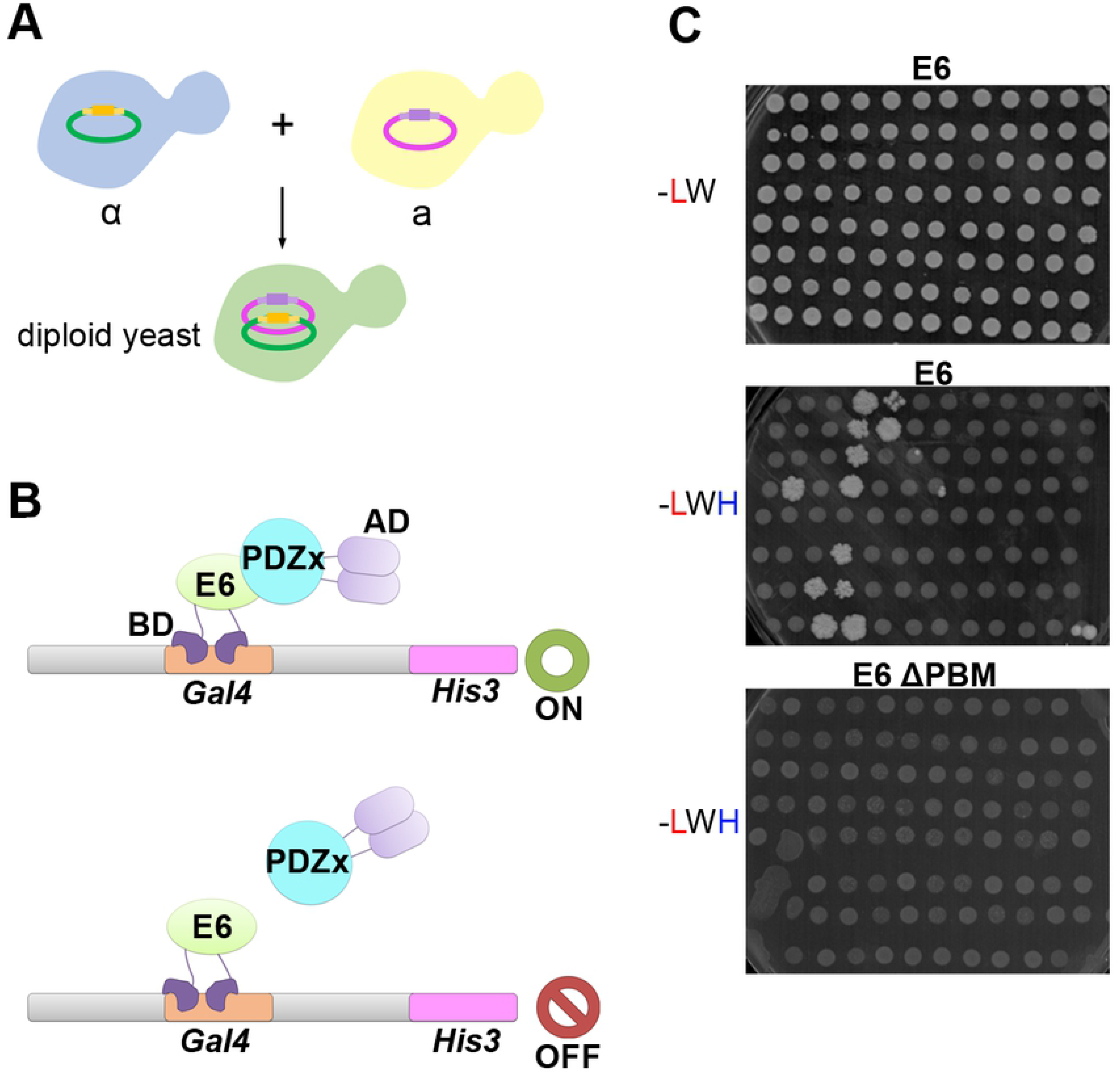
Y2H mating and selection process. **(A)** Scheme illustrating the mating of the two yeasts strains. The ‘a’ type yeasts hosting the E6-pGBT9-BD baits and the ‘α’ type yeasts hosting the PDZome 2.0-pACT2-AD were allowed to mate. Diploid yeasts containing both the PDZ and the E6 constructs were selected in synthetic agar medium. **(B)** Scheme illustrating the detection of protein interaction by Y2H. In case the E6-bait coupled to the Gal4 binding domain (BD) interacts with the given PDZ-prey coupled to the Gal4 activation domain (AD), the *HIS3* reporter gene is expressed, allowing growth of the diploid yeasts in a synthetic medium without histidine. In case there is no interaction between bait and prey, the AD is not recruited and the *HIS3* reporter gene is not expressed. (**C**) Photographs exemplifying the growth of diploid yeasts containing both the PDZ and the E6 constructs. Diploid yeasts are selected in permissive culture medium without leucine and tryptophan (-LW). White dense colonies in the -LW medium, suggest an effective mating (upper panel). Simultaneously, the phenotypic test for interactions is performed in selective culture medium without leucine, tryptophan, and histidine (-LWH). White and dense colonies in the -LWH medium correspond to interaction pairs (middle panel). Disruption of the PBM effectively impairs the appearance of white dense colonies in the - LWH medium, confirming a PBM-mediated mode of interaction (lower panel).

We identified 53 PDZ domains interacting with the PBM of the E6 protein from the HPV16. These interactions were confirmed with the tandem constructs. In addition, the tandems identified four interactions not detected when PDZ domains are taken in isolation (Fig. 3, Fig. S1). Globally, the PDZome 2.0 outperforms the previous PDZome version that solely identified 36 PDZ domain interacting with E6 (27) (Fig. 3). Nevertheless, 8 interactions observed with the PDZome were not detected with the PDZome 2.0. In total, the PDZome 2.0 identified 43 PDZ proteins and 57 PDZ domains able to interact with the E6 protein of the HPV16. The previous version of the PDZome detected 28 PDZ proteins and 36 PDZ domains.

**Fig 3.**
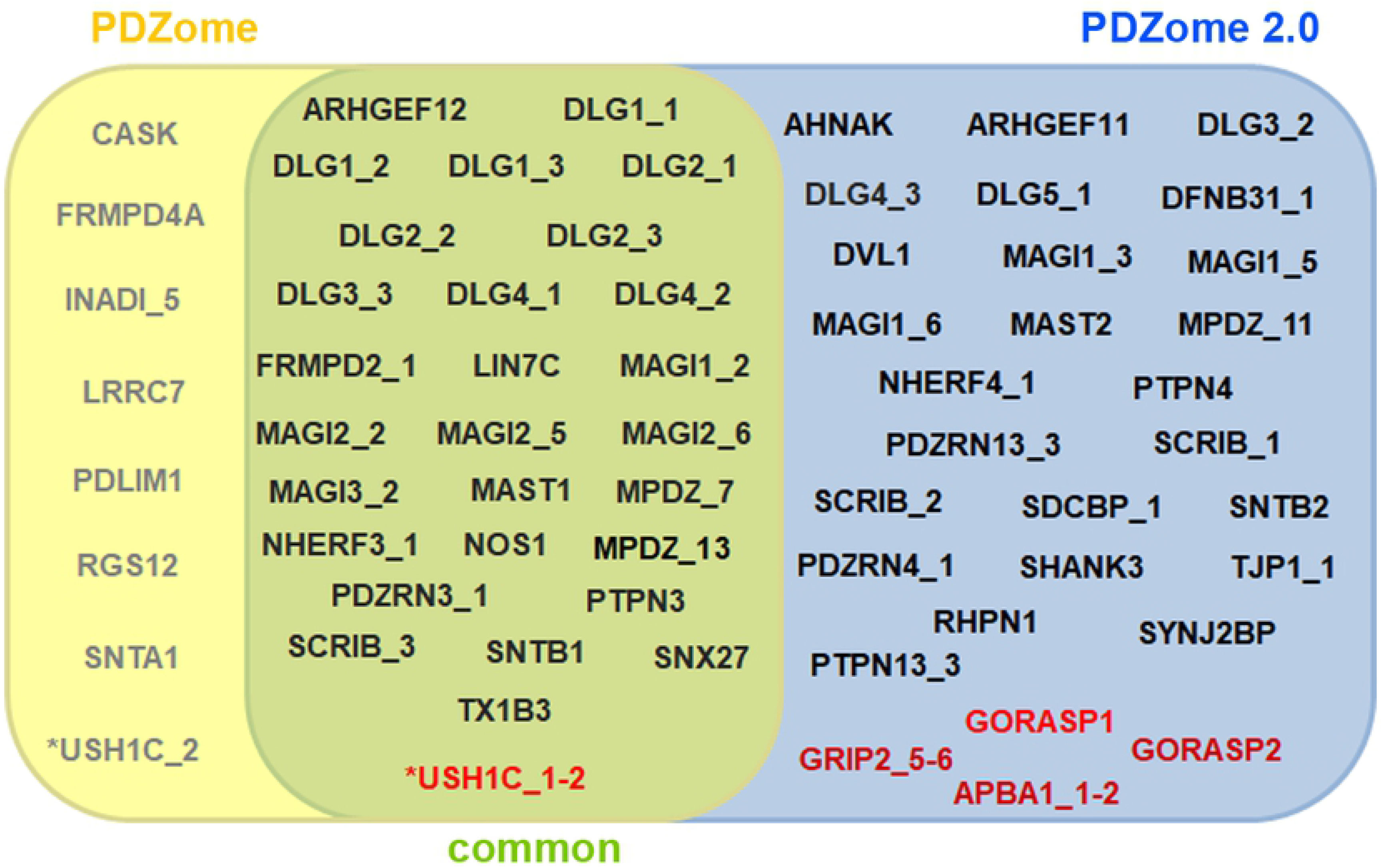
Mapping of E6-PDZ interactions using PDZome 2.0 as compared to the previous resource. Venn diagram representing the positive interactions identified by Y2H screens using the first PDZome (yellow) and PDZome 2.0 (blue). Common interactions detected using both resources are shown in the intersection region (green). Interactions revealed using PDZ tandems are highlighted in red. Note that USH1C interaction was detected using the PDZ 2 domain taken in isolation as present in the first PDZome and using the tandem (USH1C_1-2) from PDZome 2.0.

## Discussion and conclusion

In this study, we built and validated a new and most comprehensive resource to test for human PDZ interactions by Y2H. Compared to the previous version (27), this resource contains 20 additional PDZ domains (266 instead of 246). Moreover, PDZome 2.0 contains 39 PDZ domains in tandem. Finally, PDZ domains are flanked by extended boundaries meant to support proper folding (28) and avoid false negative results.

Consistently, the PDZome 2.0 revealed 25 interactions that were not detected previously for the viral oncoprotein E6. Among those 25 interactions, 9 were previously detected using the chromatographic holdup approach (HU) (38). Curiously, the PDZome 2.0 failed to detect 7 interactions that were detected with the previous version of the PDZome. The reasons are unclear. One trivial reason could be that some constructs were erroneously annotated in the PDZome compared to PDZome 2.0 (27,35,38). Another possible explanation could be that the extended sequences in the PDZ domains restrain particular interactions or contribute to the auto-inhibition of the PDZ domain (14,28). Finally, these interactions might correspond to false positives (40).

The presence of tandem structures in a protein (i.e. co-folding domains) can enhance the affinity for a particular ligand (32). Consistently, the PDZ tandem constructs not solely validated interactions observed with PDZ in isolation but also revealed additional interactions. Three of these extra interactions were not described previously in papers reporting the HPV16-E6 -PDZ interactomes (27,38,39,41–44). Obviously, the PDZome 2.0 might still be prone to false negative. It is always recommended to verify interactomes using complementary biochemical or biophysical methods such as HU or surface plasmon resonance, before performing functional analyses (38,45,46).

In conclusion, PDZome 2.0 represents a valuable additional resource to test for PDZ interactions by Y2H and certainly an easy going first line choice when one aims to investigate in a comprehensive manner the PDZ interactome of a protein of interest.

## Abbreviations

HPV16: human papilloma virus-16
PDZ: postsynaptic density-95, disc large-1, zona occludens-1
PBM: PDZ binding motif
PDZome: PDZ proteome
Y2H: Yeast two-hybrid

## Acknowledgements

This work was supported by grants from the French National Research Agency (ANR-18-CE13-0017, Project SynTEV), the Fund for Scientific Research–Flanders (Fonds Wetenschappelijk Onderzoek—Vlaanderen Grants G.0846.15 and G0C5718N), and the Institut National du Cancer (Projets Libres de Recherche “Biologie et Sciences du Cancer” INCa 9474). J.-P.B. is a scholar of Institut Universitaire de France. M.C. benefited of a CONACYT fellowship.

## Supporting information

**Fig. S1. Summary of the Yeast-Two-Hybrid raw data.** Summary of the PDZ interactions detected by Yeast-Two-Hybrid using the E6 protein C-terminal region of HPV16 wild type as bait. (**A**) Comparison of the interactions detected using the PDZome and the single PDZ collection of the PDZome 2.0 (as indicated). (**B**) Comparison of the interactions detected using the collection of tandems of the PDZome 2.0 and the correspondent PDZ taken in isolation in PDZome and PDZome 2.0 (as indicated). No interaction detected is indicated in red, positive interaction detected is indicated in blue. Numbers correspond to positive colonies **/** detected from *n* independent experiments.

**S1 Table. Single PDZ domain constructs used to comprehensively map PDZ interactions.**

**S2 Table. Tandem PDZ constructs used as preys to comprehensively map PDZ interactions.**

